# Performance of aerial *Bacillus thuringiensis* var. *israelensis* applications in mixed saltmarsh-mangrove systems and use of affordable unmanned aerial systems to identify problematic levels of canopy cover

**DOI:** 10.1101/2020.05.10.087411

**Authors:** Brian J. Johnson, Russell Manby, Gregor J. Devine

## Abstract

**BACKGROUND:** In the Australian southeast, the saltmarsh mosquito *Aedes vigilax* (Skuse) is the focus of area-wide larviciding campaigns employing the biological agent *Bacillus thuringiensis* var. *israelensis (Bti)*. Although generally effective, frequent inundating tides and considerable mangrove cover can make control challenging. Here, we describe the efficacy and persistence of an aqueous *Bti* suspension (potency: 1200 International Toxic Units; strain AM65-52) within a mixed saltmarsh-mangrove system and the use of affordable unmanned aerial systems (UAS) to identify and map problematic levels of mangrove canopy cover.

**RESULTS:** High mangrove canopy density (>40% cover) reduced product deposition by 74.5% (0.013± 0.002 μl/cm^2^ vs. 0.051± 0.006 μl/cm^2^), larval mortality by 27.7% (60.7± 4.1% vs. 84.0± 2.4%), and ground level *Bti* concentrations by 32.03% (1144 ± 462.6 vs. 1683 ± 447.8 spores ml^−1^) relative to open saltmarsh. Persistence of product post-application was found to be low (80.6% loss at 6 h) resulting in negligible additional losses to tidal inundation 24 h post-application. UAS surveys accurately identified areas of high mangrove cover using both standard and multispectral imagery, although derived index values for this vegetation class were only moderately correlated with ground measurements (*R*^2^=0.17-0.38) at their most informative scales.

**CONCLUSION:** These findings highlight the complex operational challenges that affect coastal mosquito control in heterogeneous environments. The problem is exacerbated by continued mangrove transgression into saltmarsh habitat in the region. Emerging UAS technology can help operators optimize treatments by accurately identifying and mapping challenging canopy cover using both standard and multispectral imaging.

## 1 INTRODUCTION

Control of mosquitoes in coastal saltmarsh and mangrove habitats of Australia is a major public health and economic concern due to the biting nuisance they present and the status of some species as significant vectors of Ross River (RRV) and Barmah Forest viruses (BFV).^1–5^ Broad scale control programs, run by local government and largely reliant on aerial larviciding, are the preferred form of control. The primary agent used in these campaigns is the microbial insecticide *Bacillus thuringiensis* var. *israelensis (Bti)*, a gram positive entomopathogenic bacterium, with high specificity, rapid impact and minimal ecological and environmental influence.^6–8^ These programs are associated with a reduction in the incidence of RRV^9^ making them a critical component of the regional public health program. The efficacy of these measures is, however, affected by a number of environmental and biological factors^10, 11^ and it is of fundamental importance that operators understand the impacts of such factors on this vital vector control tool.

The presence of vegetative canopy is a principal barrier to control as it reduces product deposition at ground level.^12, 13^ In coastal zones of southeast Queensland, mangroves provide canopy cover over large areas of productive mosquito breeding habitat and are the primary barrier to effective larvicide coverage in the region. Mangrove cover has intensified in recent decades in response to sea level rise and land use change.^14–16^ In some areas, saltmarsh loss has exceeded 70% due mangrove transgression. Saltmarsh mosquito densities remain high because mangroves, particularly mangrove hummocks and the water they retain, provides ample breeding habitat for primary species like *Aedes vigilax*.^17–19^ In response, many operators have opted to use higher density granular *Bti* products rather than liquid formulations to increase penetration through the canopy and improve control. Although effective,^20^ granular products often incur greater per hectare costs than liquids due to poorer logistics (i.e., moving and storing bulk product), payload disadvantages (i.e., decreased amount of treated area per sortie) and less uniform ground coverage.^21^ Thus, many operators prefer to use liquid *Bti* in open landscapes and along the saltmarsh-mangrove interface as these areas lack contiguous mangrove cover. Upwards of 50,000 L of liquid *Bti* are applied annually over approximately 40,000 ha of saltmarsh and mixed mangrove-saltmarsh habitat in southeast Queensland.^22^ Reliance on liquid *Bti*, continued landward encroachment by mangroves and the importance of the regional mosquito control program require that we understand how *Bti* products perform in changing landscapes.

Factors that decrease the active lifespan, or persistence, of *Bti* once in the aquatic mosquito habitat also impact control. Exposure to ultraviolet (UV) radiation,^23, 24^ application in organically enriched habitats,^25, 26^ sedimentation,^27, 28^ and agitation^27^ are all known to negatively affect persistence. Additional factors such as tidal diffusion or flushing are poorly understood. In the bays around southeast Queensland, islands comprise a significant percentage of saltmarsh mosquito habitat. These islands experience extended tidal inundation events that trigger large-scale synchronous egg hatch of *Ae. vigilax* and *Culex sitiens* species. During spring tides, the majority of each island is inundated for up to 4 consecutive days, with about 7% of tides resulting in complete inundation.^29^ The persistence of *Bti* is presumably minimal under these circumstances particularly for those formulations directed at single hatching events. This is supported by comparisons of persistence between areas exposed to frequent tidal inundation^30^ and static systems such as lakes^31^ and containers.^32^ Understanding and predicting the loss of control associated with tidal inundation is therefore critical to the coordination of larviciding activities.

Effective area-wide larviciding also relies upon our capacity to identify, quantify and map mosquito productivity in complex environments. The majority of operators currently rely upon satellite imagery such as Landsat and SPOT for such tasks.^33–35^ These types of medium-resolution (5-30 m) imagery provide broad-scale habitat characterizations, but they do not offer the resolution necessary for characterizing irregular canopy cover at operationally relevant scales. Although the use of advanced multispectral satellite imagery such as that obtained by Sentinel-2 and MODIS greatly improves canopy classification,^36–38^ the spatial resolution is still too poor (10-20 m Sentinel-2; >100 m MODIS) for most operational needs. The recent proliferation of unmanned aerial systems (UAS; commonly referred to as drones) now provides operators the means of obtaining very high resolution (<10 cm/pixel) imagery at relatively low cost. UASs are now routinely used to characterize canopy cover in a wide array of forest and agricultural systems.^39–42^ In mangrove systems, UAS-based predictions of leaf area cover are comparable to those from satellite imagery (WorldView-2),^43^ but have the additional advantage of being flown at low operating altitudes to avoid cloud cover and at times most convenient to field teams with limited resources. Until recently, UAS imaging has been challenged by the high cost of multispectral or hyperspectral imaging systems, restrictive regulations and costly pilot licensing,^44^ but unrestricted, consumer-level UASs (<2 kg total weight; i.e. DJI Mavic 2) are now available. Combined with a proliferation of affordable aftermarket multispectral cameras, such systems represent powerful mapping solutions that are currently be adopted to map mosquito larval habitat.^45, 46^ These developments make an investigation of their utility in other vector control settings a priority.

Here, we assessed the efficacy and persistence of a widely used aqueous *Bti* formulation, in an area of mixed mangrove-saltmarsh habitat that experiences frequent tidal inundation events. Based on patterns of droplet deposition, larval mortality and the quantification of *Bti* spores in water samples, we provide recommendations regarding the operational challenges of using these kinds of products for broad-scale control programs in coastal landscapes. We describe how the use of a low-cost consumer UAS, incorporating standard and multispectral imagery, can help predict operational efficacy under differing levels of mangrove canopy cover.

## 2 MATERIALS AND METHODS

### 2.1 Study site and sampling scheme

The study was performed on Pannikin Island (27°39’43.4“S 153°19’54.9”E), an uninhabited island located in Moreton Bay, a large, subtropical, estuarine ecosystem in southeast Queensland, Australia (Fig. 1 A). The island is characterized by three habitats: open saltmarsh (primarily *Sporobolus virginicusis* and *Sarcocornia quinqueflora*), mid-intertidal mangrove (primarily *Avicennia marina* var. *australasica*), and lower-intertidal mudflats. There is a large degree of habitat mixing, particularly on the southern half of the island. The island contains 135 ha of productive saltmarsh mosquito habitat that routinely produces vast numbers of the saltmarsh mosquitoes *Ae. vigilax* and *Culex sitiens*. Tides in the area are semidiurnal ranging from 0.8 to 2.0 m within a full monthly tidal cycle. Fringe mudflats and mangroves are inundated for 4-5 hours on a daily basis and most of the island is flooded for up to four consecutive days during spring tides. As receding tides drain water from the saltmarsh, water remains in depressions in the substratum, forming stagnant pools that persist for 1 to 7 d.^29, 47^

**Figure 1.**
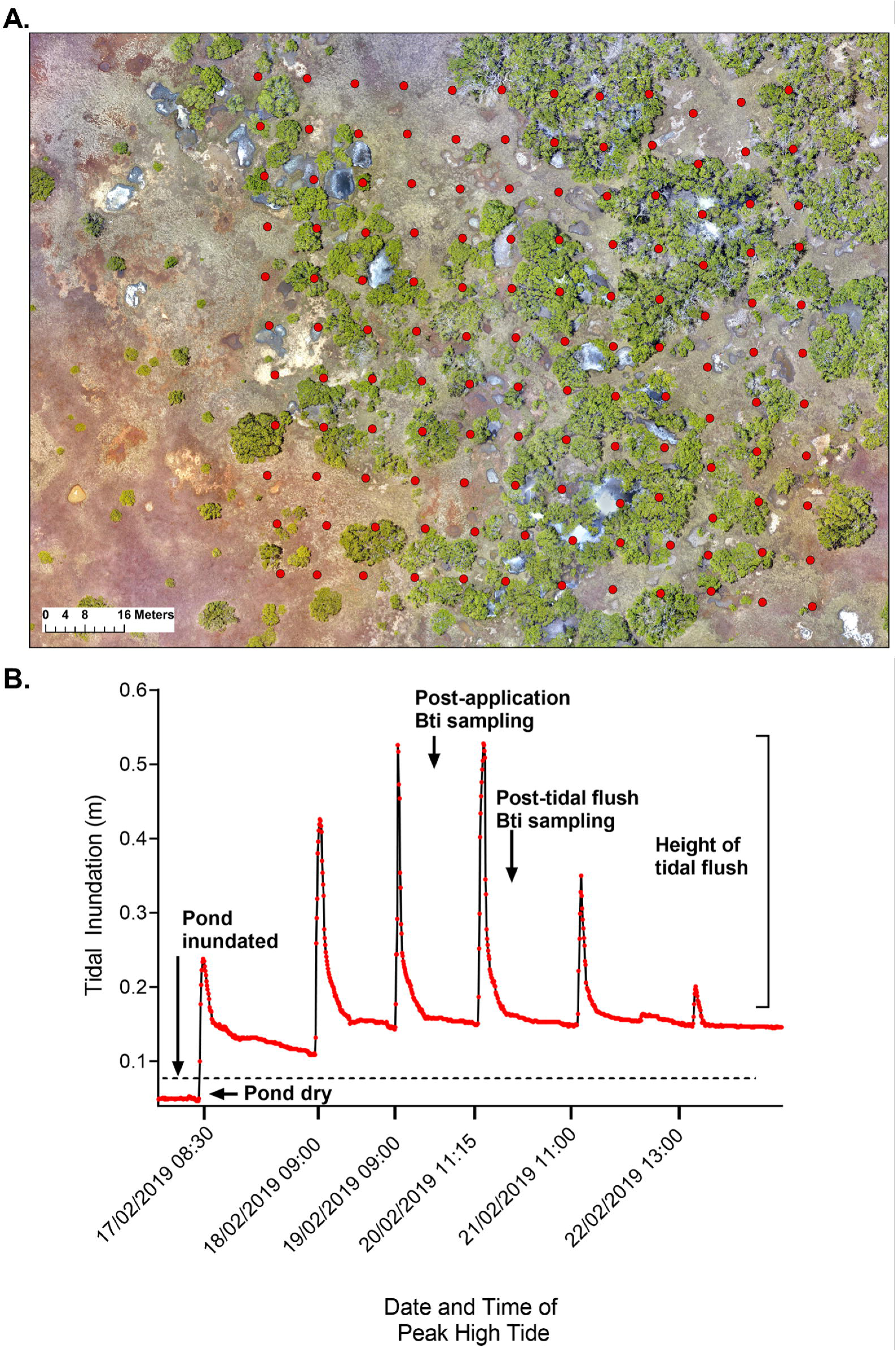
(A) Defined sampling area and location of ground sampling points (red dots) on Pannikin Island highlighting the mixture of open saltmarsh and mangrove land cover common throughout the island. (B) Tidal inundation patterns on Pannikin Island over the spring tide (‘king tide’) event studied and timing of sampling events. The figure highlights multiple significant tidal flushing events (15-46.7 cm in height) after inundation of pond features.

To capture the habitat heterogeneity on the island, we established a study area of 110 x 100 m at the junction of the saltmarsh and mangrove zones. Within that area, we established 12 sample transects each containing 11 sampling points (10 m between points, 132 sampling points in total). We monitored the timing and magnitude of daily tidal events using a Waterwatch^®^ remote water level sensor (LS1 model, Tussock Innovation, Dunedin, New Zealand) located adjacent to the sample area (Fig. 1 B).

### 2.2 Canopy cover quantification

Mangrove canopy cover at each sample point was determined by digital image analysis using the free-ware program ImageJ.^48^ Images were acquired with a digital camera (Olympus Tough TG-5) at a ratio of 4: 3 (4000 x 3000 pixels) and a focal length of 4.5 mm (aperture = f/2.0). All photos were acquired in full-auto mode. The camera was mounted horizontally on a folding tripod at a height of 1 m above the ground and levelled before use. Images were recorded under conditions of diffuse skylight during the early morning and each image was converted to white (open; sky) and black (closed; canopy) pixels. Canopy cover was then calculated as the percentage of total black pixels.

### 2.3 *Bti* formulation tested

We selected to test a widely available liquid formulation of *Bti* (Vectobac^®^ 12 AS, Valent BioSciences, Lebertyville, IL, USA; EPA Reg. No. 730449-38).^55^ The chosen product is an aqueous suspension containing crystals of *Bti* protein and bacterial spores (strain AM65-52) with a potency of 1200 International Toxin Units (ITU) per milligram (5 × 10^9^ spores ml^−1^). The active *Bti* component (fermentation solids and solubles) constitutes 11.61% of the end commercial product with the rest (88.39%) consisting of non-active ingredients. Proxel GXL^®^ (Lonza, Group LTD, Basel, Switzerland; dipropylene glycol solution of 1,2-benzisothiazolin-3-one) comprises 0.10% of the non-active ingredients, with the rest being withheld as trade secret.

### 2.4 Aerial application parameters and deposition characterization

Product deposition in areas of open saltmarsh and under mangrove canopy was determined using water-sensitive photographic paper. A single card (76 x 52 mm, 250 gsm, high-gloss photo paper, Krisp, Hoppers Crossing, VIC, Australia) was positioned horizontally 1 m off the ground by fixing the card to a wooden stake with a 22 mm bulldog clip. The chosen *Bti* formulation was applied over the study area as part of a routine larviciding operation by a Bell 47 helicopter flying at 55 knots. Product was applied at 1.2 L/ha (maximum label rate) with a target droplet size of 400 µm controlled through air-driven atomisers (Micronair AU7000; Micron Sprayers Ltd., Herefordshire, United Kingdom). Prior to application, a fluorescent red dye (Big Red Spray Marker Dye; Sipcam Pacific Pty Ltd., Geelong, VIC, Australia) was added to the spray solution at 50 ml/100 L to enable visualization of droplets on the cards. Cards were collected 15 min after aerial deposition to allow droplets to settle and stored until analysis. Droplet characterization and spray coverage was determined using the DepositScan^49^ plugin for ImageJ on scanned (600 dpi) images.

### 2.5 Larval bioassays

Expected mortality at each sampling point was estimated using larval bioassays. A single 500 ml plastic cup (10 cm diameter) was set at each sample location to collect *Bti* droplets. The cups were positioned 1 m off the ground to avoid any interference from ground vegetation. The bioassay cups were affixed vertically to the same wooden stakes used to support the droplet cards using bulldog clips. All cups were collected at least 15 min after product application and stored on ice for transport to the laboratory on the same day. Once at the laboratory, 500 ml of fresh seawater was added to each cup with efforts made to ensure all visible droplets from the sides of the containers were washed to the center. Twenty field-collected, 2^nd^ instar *Ae. vigilax* larvae were then placed into each cup with a pinch of crushed Lucerne pellet as a food source. Mortality was recorded at 24 h and 48 h.

### 2.6 Water sampling and *Bti* quantification

The concentration of *Bti* (spores ml^−1^) in ground water before and after aerial application and following a subsequent tidal flush (43.8 cm; 20 h post-application) was determined by real-time quantitative PCR (qPCR) analysis of 200 ml ground water samples collected from a subset of sampling points. Post-application water samples were collected one hour after application to allow the product to settle. A total of 60 water samples were collected, 20 for each canopy classification (i.e., high, low and open). An additional 20 pre-application control samples were collected 2 h before application and prior to peak high tide. After collection, all water samples were sealed, stored on ice and transported to the laboratory where they were stored at −20 C until processing. To investigate the impact of tidal flushing on the persistence of *Bti*, an additional 40 water samples were taken 2 h after the following high tide (20 h post application). Immediately prior to product application, water quality parameters were measured at 45 sampling points across the site using a Hanna HI9829 multi-parameter meter (Hanna Instruments, Keysborough, VIC, Australia). The measured variables included water temperature, pH, dissolved oxygen (% and ppm), salinity and turbidity (formazin nephelometric units; FNU).

Following the initial field survey, an additional persistence study was performed to better understand the persistence of *Bti* in the absence of tidal flushing. Sampling was performed during a minor tide event (incomplete inundation) following a 5 month period when no aerial larviciding operations had taken place ensuring remnant *Bti* was minimal. Liquid *Bti* was applied aerially as described above at a secondary site with similar ground vegetation characteristics as Pannikin Island. The change in site was due to logistical challenges at the time. The quantity of *Bti* spores was determined at 1 h, 6 h and 20 h post-application by taking ten 200 ml water samples at each time point. The samples were collected, stored and processed as described above.

The number of *Bti* spores in water samples was determined following standard protocols.^50, 51^ To create a qPCR standard curve, stock *Bti* solution was serially diluted in a total volume of 200 ml of saltwater to obtain a range of 1×10^8^ to 1×10^0^ spores ml^−1^. The initial spore concentration of the stock solution was determined by colony-forming unit (CFU) quantification of ten-fold dilutions on blood agar media (trypticase soy agar with 5% sheep blood, Edwards Group Pty., Narellan, NSW, Australia). CFU quantification revealed the stock concentration to be 1.20 × 10^9^ spores ml^−1^. Prior to DNA extraction, water samples were filtered through 0.45 µm mixed cellulose ester filters (Merck, Darmstadt, Germany) to concentrate spores. The filtered material was resuspended in 1 ml of distilled sterile water by briefly vortexing and left to stand for 15 min at room temperature. Resuspensions were stored at −20 C until processed. DNA was extracted from 400 µl of that resuspended material following a standard protocol^51^ with the exception that we used the DNeasy Blood and Tissue kit from Qiagen (Valencia, CA, USA) to extract DNA from sample aliquots. DNA extraction followed the specified DNeasy protocol for gram-positive bacteria. A standard curve was generated on the prepared standards using recommended TaqMan qPCR assay and thermal cycling conditions.^50^ All samples were tested in triplicate on three separate assay runs for a total of 9 replicates for each dilution. Quantitative PCR was carried out in duplicate for each field collected sample using the same DNA extraction and PCR conditions.

### 2.7 UAS surveying and image processing

Three separate UAS surveys were carried out in February 2018. The first survey was performed using a DJI Phantom 4 (DJI, Shenzhen, China) quadcopter system fitted with a 4K camera (7.81 mm CMOS sensor, 12 Megapixel) with an f/2.8 lens and a 94° field of view. The second survey used a DJI Mavic Pro (DJI, Shenzhen, China) quadcopter also fitted with a 4K camera (4K, 12 MP, 2.3” CMOS sensor, 12 Megapixel) with an f/2.2 lens and a 78.8° field of view. During each of these surveys, the UASs were operated at an altitude of 40 m in a double grid pattern with 80% front overlap and 70% side overlap. This resulted in a ground sampling distance (GSD) of <2 cm/pixel. A third survey was performed using a DJI Mavic Pro fitted with a multispectral camera (red-green-near infrared; MAPIR^®^ Survey 3; 12 MP; San Diego, CA) at the same flight parameters as the previous surveys. Flights were controlled through the Pix4D mobile application. Captured RGB images were processed using the Pix4Dmapper and Pix4Dfields applications to generate site orthomosaics and Visible Atmospherically Resistant Index (VARI) and Triangular Greenness Index (TGI) outputs. The multispectral images captured by the MAPIR^®^ camera were pre-processed and calibrated to a reference reflective target (MAPIR^®^ Calibration Target V2) using the MAPIR^®^ provided software and workflow. Calibrated images were then uploaded and a site orthomosaic created in Pix4DMapper. The generated georeferenced tiff file was imported into ArcGIS Pro (ESRI, Redlands, CA) to generate an NDVI image using the Image Analysis workflow. Supervised (object-based) image classification was then performed using ground-truthed vegetation and canopy cover plots to train the image classifier (Support Vector Machine; 500 samples per class). Image classification on the RGB-based indices provided poor separation of identified vegetation classes due to high value overlap. Instead, high-density cover was estimated by extracting the total number of pixels falling within the 95% CIs of the mean index value (zonal distance = 2 m) for high canopy cover (>40%). Low-density estimates were not attempted due to the high overlap in index values between classes. The upper limit of the TGI calculation was extended to the maximum index value as the use of only the upper 95% CI greatly underestimated canopy cover. Additional zonal summaries were generated at distances of 1 m, 5 m and 10 m from the center of each ground sample point for regression analysis against ground truthed canopy measurements.

### 2.8 Statistical methods

Preliminary investigations revealed that three distinct canopy cover classifications existed across the study site. These included areas of high (>40%) and low (<40%) canopy cover and open saltmarsh areas. Differences in product deposition, spray characteristics (droplet size, density, and coverage), larval mortality and *Bti* concentrations across the three canopy classifications were compared using nonparametric Kruskal–Wallis tests to address unequal sample sizes across the three classifications. All datasets were found to be non-normally distributed and therefore percent mortality data was arc-sine transformed prior to analysis. Differences in the concentration of *Bti* before and after aerial application and following an ensuing tidal flush event were analyzed by Kruskal-Wallis tests to address data that violated assumptions of normality and homogeneity of variances. Standard Major Axis (SMA) regression analysis was used to analyze the relationship between ground-level canopy cover estimates and generated zonal summaries for each vegetation index. SMA regression analysis accounts for uncertainties in both sets of measurements and allows testing the statistical significance of any deviation of the slope from unity and the intercept from zero.^52^ All statistics were performed using the R^53^ and Prism (version 8.1.2; GraphPad, San Diego, CA) statistical packages.

## 3 RESULTS

### 3.1 Water quality

No significant differences (*F*_*2, 352*_= 0.75, *P*=0.48) in measured water quality variables was observed across the study site (Table S1). All values were remarkably similar across the three canopy classifications with the exception of greater turbidity observed in high canopy areas (21.5± FNU vs. 11.6± and 12.7± FNU) and slightly greater water temperatures measured in open saltmarsh areas relative to those under low and high canopy cover (35.9 C vs. 34.9 C and 34.9 C).

### 3.2 Canopy cover and its impact on spray penetration and larval mortality

Canopy cover density (percent canopy) ranged from 10-77% (*n=*55; mean = 40.3%; 95% CI: 34.9-45.7%). Application in areas of high canopy density (*n=*35) significantly affected all droplet characteristics except for mean volumetric diameter (0.5 VDM; Table 1). Overall, product deposition was reduced by 75.2% in areas of high canopy density (Fig. 2 A) relative to open areas (0.01± 0.002 µl/cm^2^ vs. 0.05± 0.006 µl/cm^2^) and by 60.7% relative to areas of low canopy cover (*n=*20, 0.03± 0.007 µl/cm^2^). This reflected a decrease in the number of droplets reaching the ground and poorer card coverage in areas of high canopy cover.

**Table 1.**
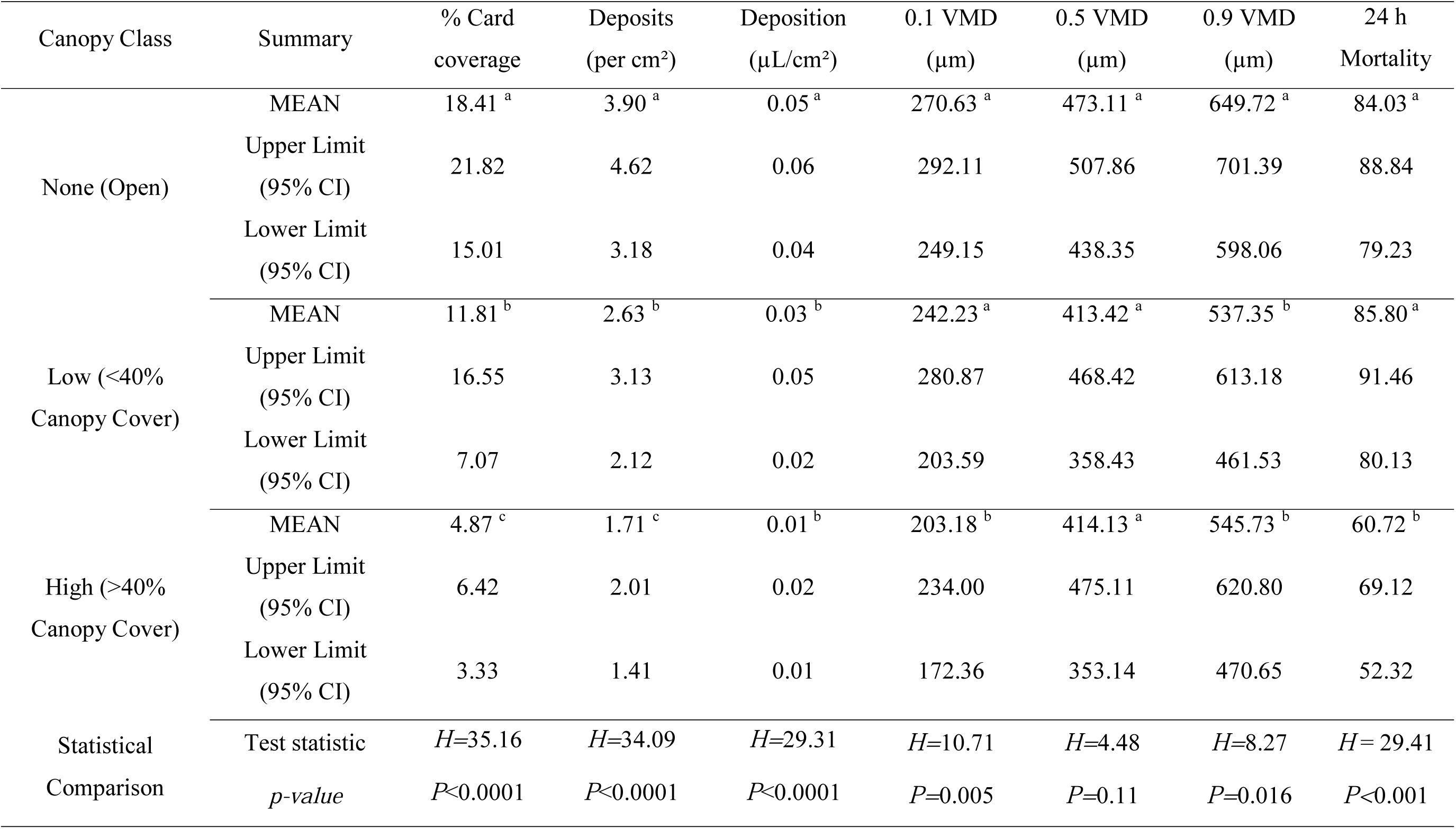
Summary of *Bti* spray characteristics by canopy cover class. Differing superscript letters within the same column represent statistically significant differences (*p-value*<0.05, Kruskal-Wallis Test).

| Canopy Class | Summary | % Card coverage | Deposits (per cm<sup>2</sup>) | Deposition (μL/cm<sup>2</sup>) | 0.1 VMD (μm) | 0.5 VMD (μm) | 0.9 VMD (μm) | 24 h Mortality |
| --- | --- | --- | --- | --- | --- | --- | --- | --- |
| None (Open) | MEAN | 18.41 <sup>a</sup> | 3.90 <sup>a</sup> | 0.05 <sup>a</sup> | 270.63 <sup>a</sup> | 473.11 <sup>a</sup> | 649.72 <sup>a</sup> | 84.03 <sup>a</sup> |
|  | Upper Limit (95% CI) | 21.82 | 4.62 | 0.06 | 292.11 | 507.86 | 701.39 | 88.84 |
|  | Lower Limit (95% CI) | 15.01 | 3.18 | 0.04 | 249.15 | 438.35 | 598.06 | 79.23 |
| Low (<40% Canopy Cover) | MEAN | 11.81 <sup>b</sup> | 2.63 <sup>b</sup> | 0.03 <sup>b</sup> | 242.23 <sup>a</sup> | 413.42 <sup>a</sup> | 537.35 <sup>b</sup> | 85.80 <sup>a</sup> |
|  | Upper Limit (95% CI) | 16.55 | 3.13 | 0.05 | 280.87 | 468.42 | 613.18 | 91.46 |
|  | Lower Limit (95% CI) | 7.07 | 2.12 | 0.02 | 203.59 | 358.43 | 461.53 | 80.13 |
| High (>40% Canopy Cover) | MEAN | 4.87 <sup>c</sup> | 1.71 <sup>c</sup> | 0.01 <sup>b</sup> | 203.18 <sup>b</sup> | 414.13 <sup>a</sup> | 545.73 <sup>b</sup> | 60.72 <sup>b</sup> |
|  | Upper Limit (95% CI) | 6.42 | 2.01 | 0.02 | 234.00 | 475.11 | 620.80 | 69.12 |
|  | Lower Limit (95% CI) | 3.33 | 1.41 | 0.01 | 172.36 | 353.14 | 470.65 | 52.32 |
| Statistical Comparison | Test statistic | *H*=35.16 | *H*=34.09 | *H*=29.31 | *H*=10.71 | *H*=4.48 | *H*=8.27 | *H* = 29.41 |
|  | *p-value* | *P*<0.0001 | *P*<0.0001 | *P*<0.0001 | *P*=0.005 | *P*=0.11 | *P*=0.016 | *P*<0.001 |

**Figure 2.**
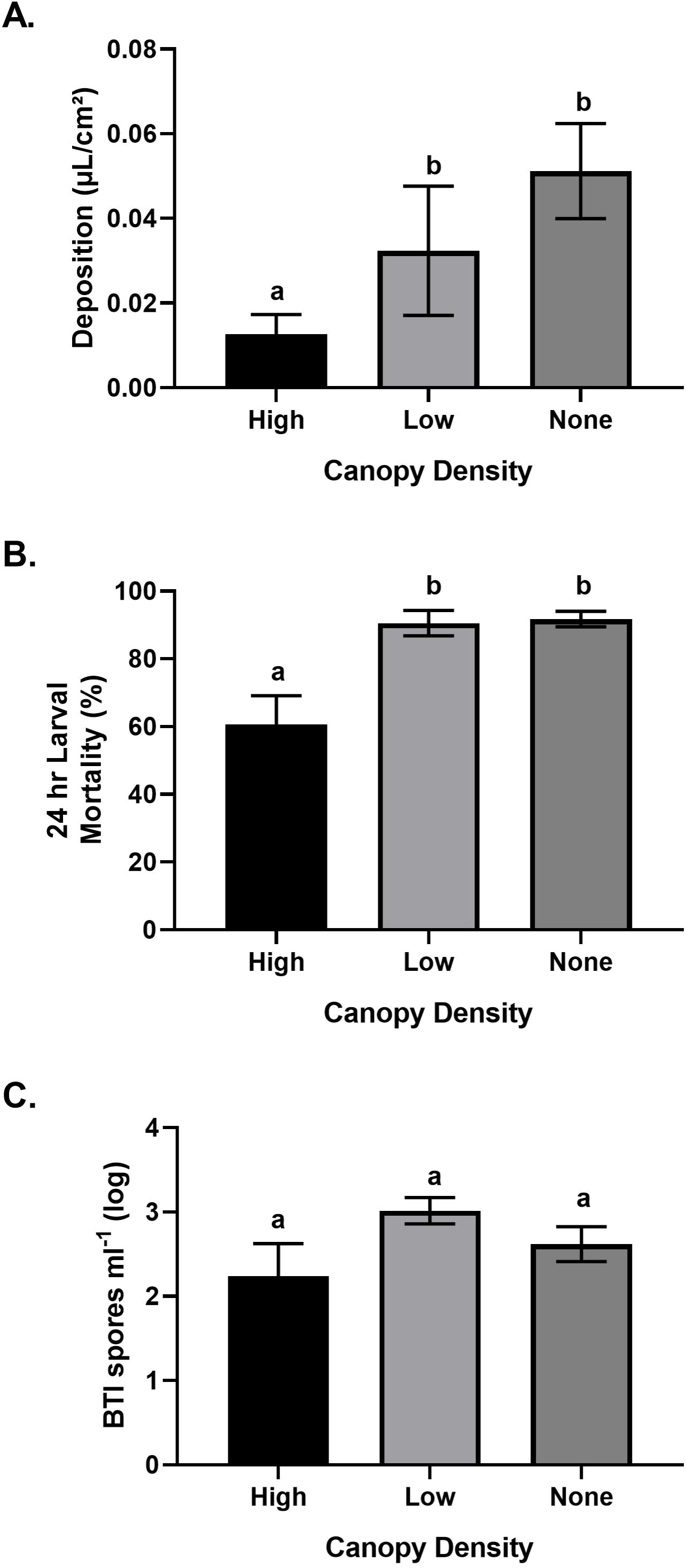
(A) *Bti* deposition (µl/cm^2^) determined from droplet card analysis, (B) 24-h larval mortality (%) observed in bioassay cups, and (C) *Bti* spore (log) concentrations (spores ml^−1^) in ground water 1 h post-application (rate= 1.2 Lt/ha) by canopy cover class. Different letters denote statistical (*p*-value<0.05) significance. Canopy density classes include high (>40%) and low (<40%) canopy cover and open saltmarsh.

Decreased product deposition in high-density canopy areas was related to a 27.7 % reduction in 24 h larval mortality (Fig. 2 B) of *Ae. vigilax* relative to open areas (60.7± 4.1% vs. 84.0± 2.4%) and 29.3% relative to areas of low canopy cover (85.8± 2.%) as revealed by larval bioassays. These reductions decreased to 24.6% (70.2± 4.6% vs. 93.2± 1.5%) and 24.2% (70.2± 4.6% vs. 92.7± 2.4%), respectively, at 48 h.

### 3.3 Impact of canopy density on ground-level *Bti* concentrations

Of the 60 water samples collected post application, 49 produced a successful qPCR result resulting in uneven samples sizes for each canopy density classification (open= 26; low=12; high=11). Overall, there was a 36.8% and 32.0% reduction in ground-level *Bti* concentrations in areas of high canopy density (Fig. 2 C; 1144 ± 462.6 spores ml^−1^) relative to those recorded under low canopy density (1811 ± 535.6 spores ml^−1^) and in open saltmarsh (1683 ± 447.8 spores ml^−1^), respectively. However, nonparametric analysis revealed no significant (*H=*1.85, *p-*value=0.40) differences in *Bti* concentrations among the canopy density classifications.

### 3.4 *Bti* persistence in the absence and presence of a flushing tide

*Bti* exhibited low persistence in the field in both the presence (peak inundation of 43.8 cm) and absence of a flushing tide (Table 2). *Bti* concentrations decreased significantly (*F_2,51_=*7.14, *p*-value=0.002) by 80.6% and 91.1% at 6 h (294.3±77.3 spores ml^−1^) and 24 h (135.5±27.7 spores ml^−1^) post-application, respectively, relative to concentrations at 1 h post-application (1515±364.9 spores ml^−1^). A similar loss of *Bti* was observed (97.1%; 1498± 321.6 vs. 43.3± 9.9 spores ml^−1^) following a single flushing tide event 20 h post-application.

**Table 2.** *Bacillus thuringiensis* var. *israelensis* concentrations (spores ml^−1^) before and after aerial application (product application rate= 1.2 Lt/ha) in the absence and presence of a flushing tide event.

| Sampling Point | Persistence: No flushing tide |  |  |  | Persistence: flushing tide (20 h post-treatment) |  |
| --- | --- | --- | --- | --- | --- | --- |
|  | Pre-treatment<br>Control | 1 h Post<br>Treatment | 6 h Post<br>Treatment | 24 h Post<br>Treatment | 1 h Post<br>Treatment | Post Tidal<br>Flush |
| Number of values | 18 | 9 | 9 | 9 | 33 | 22 |
| Mean | 14.77 | 1515 | 294.3 | 135.5 | 1498 | 43.25 |
| Std. Error of Mean | 4.436 | 364.9 | 77.31 | 27.7 | 321.6 | 9.999 |
| Lower 95% CI | 5.365 | 744.7 | 131.2 | 77.04 | 843.3 | 22.46 |
| Upper 95% CI | 24.17 | 2285 | 457.4 | 193.9 | 2153 | 64.04 |
| Percent (%) loss (relative to<br>1 h post-treatment) | N/A | N/A | 80.57 | 91.06 | N/A | 97.11 |

### 3.5 UAS imaging to map problematic levels of mangrove canopy density

Both RGB-based indices readily separated areas of mangrove canopy cover from other land cover classes; however, VARI misrepresented areas of water as vegetated areas to a much greater degree than TGI (Fig. 3 A, B). This misrepresentation was corrected when isolating index values associated with high-canopy density (Fig. 4 A, B), whereas TGI differentiated these areas correctly without isolation of canopy index values. Overall, each index produced modest correlations with ground level canopy measurements across all zonal distances (1, 2, 5 and 10 m; Fig. 5 A-H). The strongest correlations were observed for each index at a zonal distance of 5 m with VARI (*R*^2^=0.38; *P*<0.001) outperforming TGI (*R*^2^=0.17; *P*<0.02). A total of ca. 23.5% of the imaged area (0.35 ha out of 1.5 ha) was determined to be covered by high-density canopy using VARI and 36.6% (0.55 ha out of 1.5 ha) using TGI.

**Figure 3.**
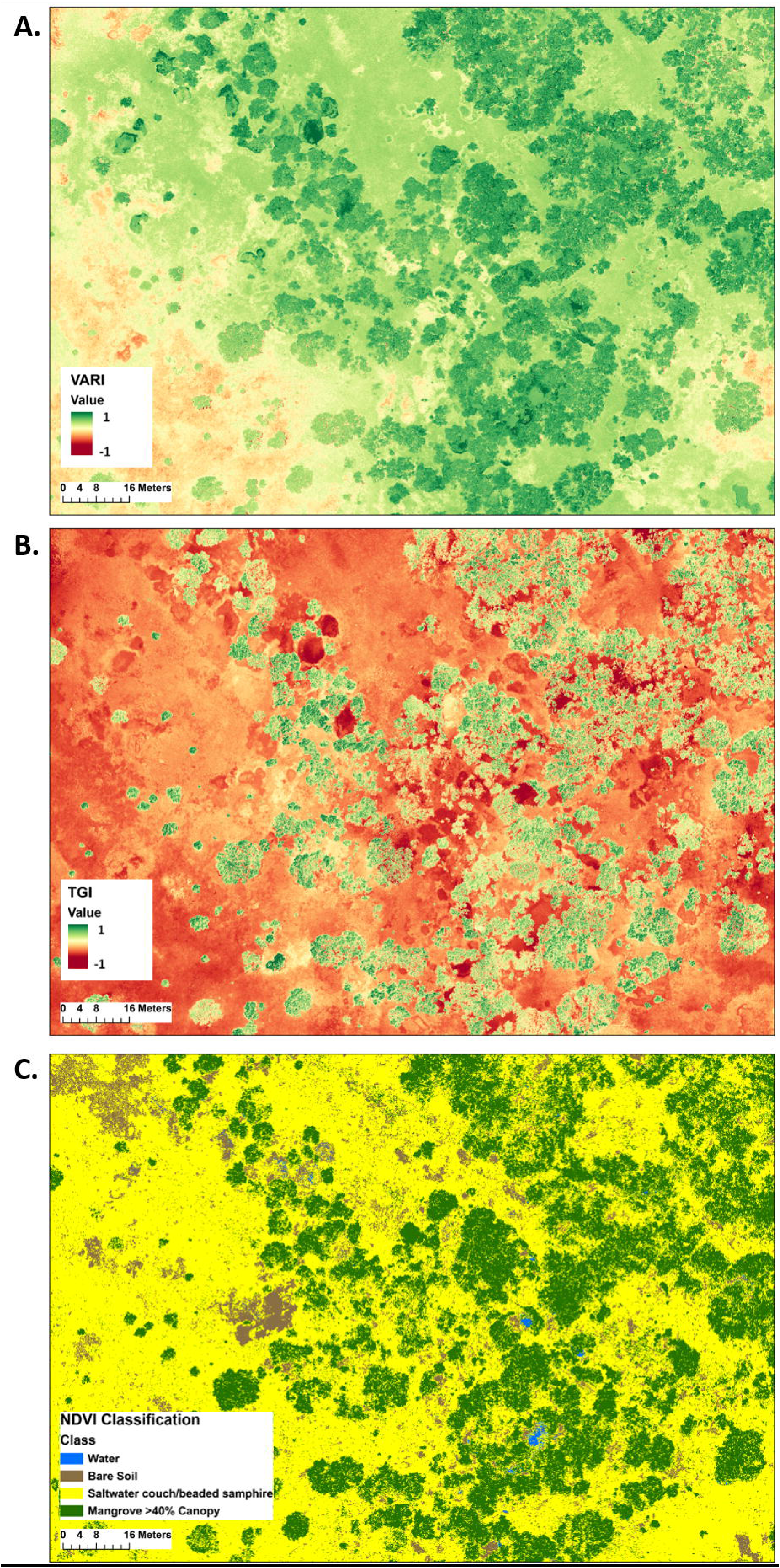
Example outputs from obtained UAS imagery, including the (A) Visible Atmospherically Resistant Index (VARI), (B) Triangular Greenness Index (TGI), and classified Normalized Difference Vegetation Index (NDVI). The area was imaged between large tide events and during a period of low rainfall.

**Figure 4.**
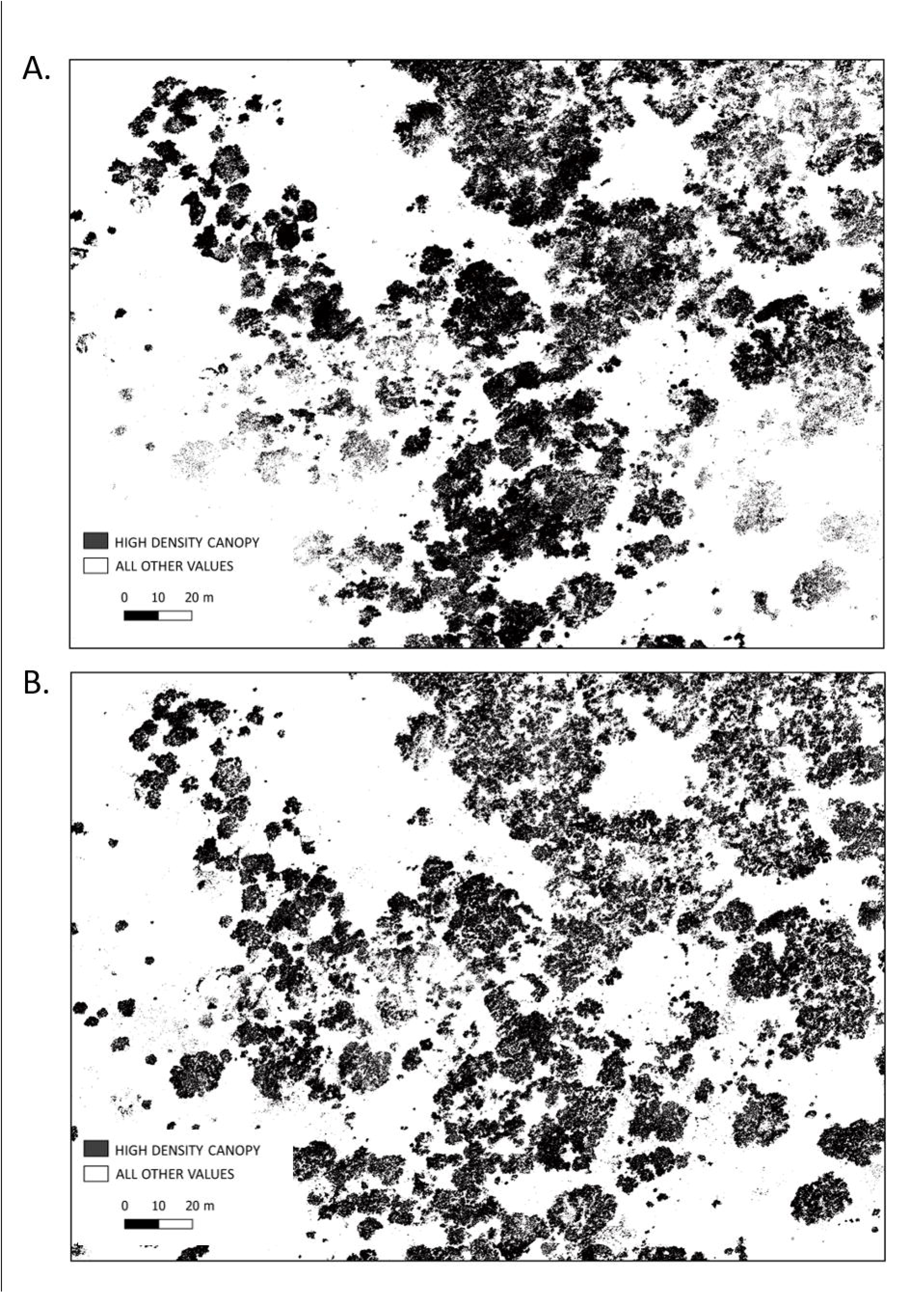
Comparison high-density canopy cover identified using the (A) Visible Atmospherically Resistant Index (VARI) and the (B) Triangular Greenness Index (TGI). Comparing the images demonstrates the slightly more conservative mapping of mangrove canopy cover by using VARI relative to the TGI.

**Figure 5.**
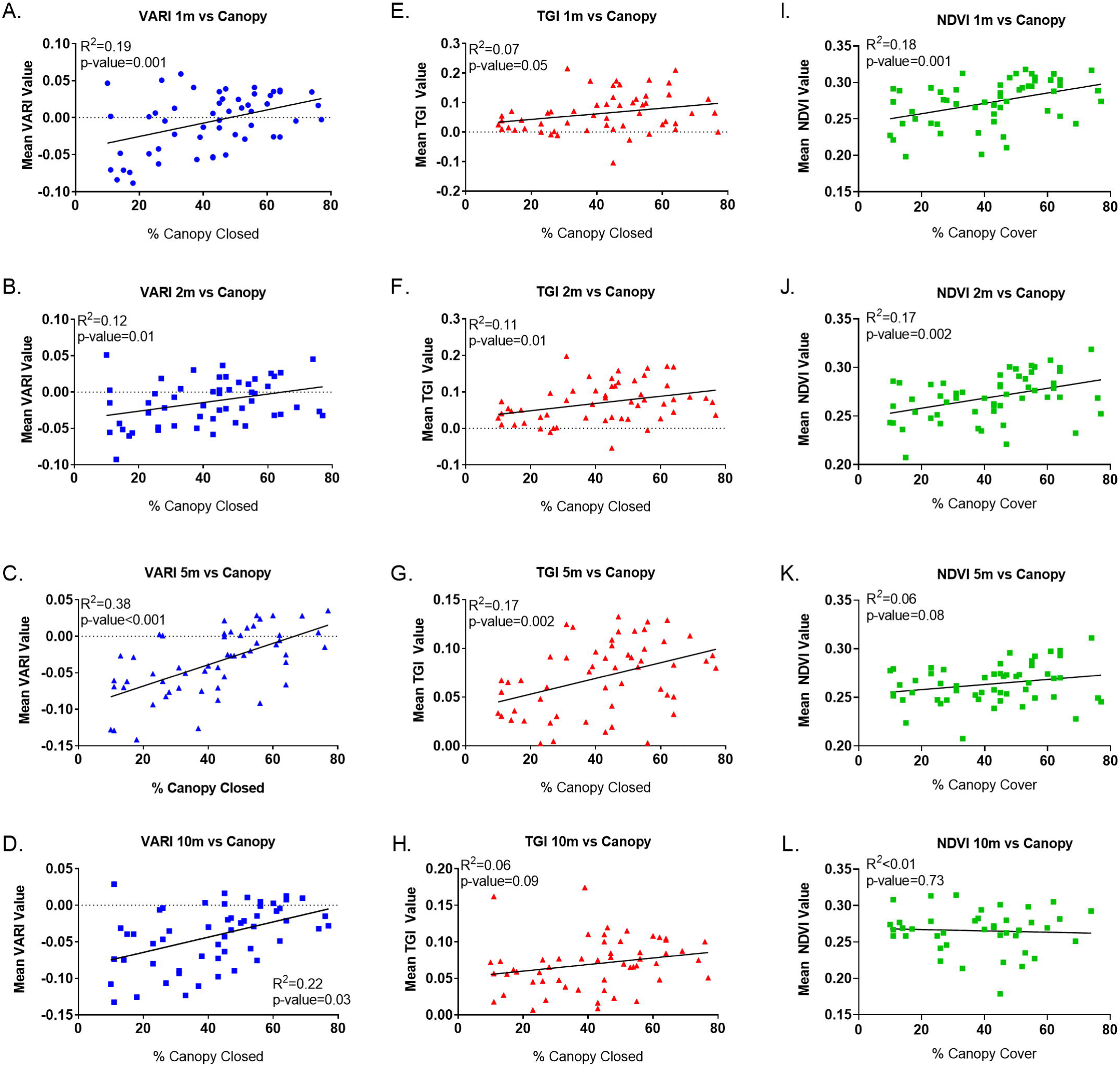
Summary of relationship between UAS acquired vegetation indices (VARI, TGI and NDVI) and ground level canopy measurements across a range of outputs.

The inclusion of multispectral imagery and resulting NDVI outputs enabled the accurate separation of primary vegetation/land classes (Fig. 3 C), although relationships with ground level canopy measurements were similar to those obtained by the RGB-based indices (Fig. 5 I-L). This allowed for accurate mapping of mangrove and saltmarsh cover and separation of bare soil from water, classes that are either incorrectly recorded as high/low canopy cover (VARI) or difficult to separate (TGI) in the RGB indices due to the high reflectance of sunlight from these surfaces. Similar to the RGB-based indices, separation of low and high-density mangrove canopy was unsuccessful due to high value overlap. NDVI classification revealed 26.5% (0.40 ha) of the imaged area to be covered by high-density canopy and 45.1% (0.68 ha) being covered by saltmarsh vegetation (i.e., saltwater couch and beaded samphire) with the rest being classified as bare soil (29.8%) or water (4.5%). Overall, the use of multispectral imagery was found to be the most advantageous for mapping exercises, whereas the TGI is preferred when multispectral imagery is not available.

## 4 DISCUSSION

*Bti* provides an effective alternative to broad spectrum larvicides and has become the principal non-chemical means of larval mosquito control in fresh and saltwater environments, yet its efficacy is strongly regulated by site-specific biotic and abiotic factors.^10^ Our study reinforces previous observations of poor penetration of liquid *Bti* formulations through tree canopy^12, 13, 54^ and low persistence in non-static systems.^30^ Despite underperformance in areas of high canopy cover, the liquid *Bti* product tested performed adequately in areas of low canopy cover and should remain a viable control tool in areas with noncontiguous or sparse mangrove canopy cover, particularly once logistical and payload advantages are considered.^21^ Emerging UAS technology can help simplify product selection and optimize control strategies by accurately identifying and mapping levels of canopy cover associated with suboptimal control using both standard and multispectral imaging. These relatively inexpensive tools provide mosquito control services critical information for directing and predicting the impacts of local operations.

Observations from this and other studies strongly suggest that operators give preference to granular products over liquids when applying in areas of moderate to high canopy density to increase product penetration at ground level.^54^ The logistical advantages (weight, cost and uniform deposition) of liquids suggests that we should continue to investigate their limitations. This includes an optimization and revision of label rates by manufacturers. Moderate increases in application rates, if beneficial, will keep the operational costs of liquids below those of granular alternatives with little inherent risk due to their limited persistence, high target specificity and minimal ecological impact at rates above those tested here.^6–8^ Strategies involving the manipulation of droplet size lack evidence of their benefit to canopy penetration and smaller droplets may reduce deposition loss due to canopy impact. The actual amount of material reaching the ground under these conditions is poorly documented.^54^ Attempts to increase deposition by using “raindrop” nozzles that produce extremely large droplets that “punch” their way through the canopy have been proposed, but this concept needs evaluation and would greatly reduce the effective swath width of the product.

The persistence of *Bti* larvicidal activity is severely curtailed in organically enriched waters like those typified by productive coastal wetlands and saltmarshes.^6, 25, 55^ Therefore, the poor persistence of *Bti* observed in this study, even in the absence of a tidal flushing event is unsurprising. Movement of product through the water column, or settling, also impacts persistence, especially against mosquito species that feed at the surface or that filter-feed rather than browse.^28^ Turbidity can also accelerate adsorption and settling and may exacerbate conditions under high canopy density that were observed to have higher turbidity. Another complicating factor is the high density of larvae encountered in coastal environments. A negative linear correlation between larval density and the efficacy of *Bti* has been observed for both *Aedes vexans* and *Culex pipiens*.^11^ This suggests that average larval densities need to be determined before the optimal dosage for a routine treatment can be established; however, such estimates are unlikely to be accurate for any given application as the density of larvae can vary greater between hatching events and at different periods of inundation. Single-brood granular products, including water dispersible granules, have similar persistence profiles relative to their liquid counterparts,^20, 56^ whereas the use of slow-release formulations may improve persistence^57^ if registered and available.

Mapping of complex larval habitats is essential to maintaining operational success and efficiency of aerial programs. The low-cost UAS tested in this study promised a useful, high resolution (<10 cm/pixel) analysis of treatment areas. Orthomosaics (RGB and multispectral) could be produced quickly and proved effective at mapping broad levels of canopy cover. Licensing costs for post-processing software (e.g. DroneDeploy, Pix4D and ArcGIS Pro) can be expensive (>$150 USD/month) but freeware and low-cost alternatives (e.g., QGIS and OpenDroneMap) are also available. Discrepancies between remotely-sensed and ground-level canopy measurements as observed in this study are comparable to those reported by others.^58^ Such discrepancies are likely attributable to canopy cover heterogeneity and how different imaging methods capture this heterogeneity at varying scales. Additional work incorporating other standard measures of canopy structure such as leaf area index^38, 43, 59^ is warranted. Comparison of selected RGB-based indices revealed that the VARI misrepresented areas of water as vegetated areas to a greater degree than the TGI, but separation of these classes is possible with additional analysis. The greater ability of the TGI to separate these classes is likely attributable to the greater sensitivity of this index to chlorophyll content.^60^ The incorporation of multispectral imaging enabled accurate mapping of the dominant vegetation classes and the accurate separation of bare soil from water. Thus, the use of multispectral imagery is preferred in mapping exercises, whereas the TGI is advantageous when multispectral imagery is not available.

## 5 CONCLUSION

As with traditional insecticides, the efficacy of *Bti* is highly contingent upon the biotic and abiotic characteristics of the environment in which it is applied. When such factors are studied and accounted for, *Bti* can used successfully in complex operational environments and is highly desirable as it is one of the few biological, ecologically harmless and largely unresisted tools available for broad scale larviciding. The emergence and proliferation of affordable UAS technology allows operators to assess and plan their activities in ever greater detail helping maintain the operational viability of available vector control tools and formulations.

## ACKNOWLEDGEMENTS

We thank the staff of the Redland City Council Pest Management team for their assistance with trapping mosquitoes and for providing transport to field sites. The Mosquito and Arbovirus Research Committee (Inc.), an independent Australian body involved in mosquito research made up of representatives from local governments, state government agencies, industry and scientific organizations, provided the funding for the study.

