## Supplementary material for "Performance of aerial *Bacillus thuringiensis* var. *israelensis* applications in mixed saltmarsh-mangrove systems and use of affordable unmanned aerial systems to identify problematic levels of canopy cover": Table S1

**Supplementary Data**

**Table S1.** Water quality values determined for a subset of sampling points distributed across the study site for each canopy classification. All measurements were taken using a Hanna HI9829 multi-parameter meter (Hanna Instruments, Keysborough, VIC, Australia).

| Canopy Classification | High Canopy Density | Low Canopy Density | Open Saltmarsh |
| --- | --- | --- | --- |
| Temp.[°C] | 34.94 | 34.90 | 35.86 |
| pH | 6.40 | 6.42 | 6.41 |
| TDS [ppm] | 27512 | 25369 | 25335 |
| Sal.[psu] | 36.12 | 33.43 | 32.94 |
| Press.[psi] | 14.47 | 14.46 | 14.46 |
| D.O.[%] | 48.78 | 46 | 54.36 |
| D.O.[ppm] | 2.85 | 2.72 | 3.17 |
| Turb. FNU | 21.5 | 11.56 | 12.66 |
